# Mechanistic basis of dynamic and heterogeneous divisive normalization in visual cortex

**DOI:** 10.1101/2025.08.19.671076

**Authors:** Harold Rockwell, Jason N. MacLean

**Affiliations:** Committee on Computational Neuroscience, University of Chicago, Chicago, IL, United States of America; Department of Neurobiology, University of Chicago, Chicago, IL, United States of America; Neuroscience Institute, University of Chicago, Chicago, IL, United States of America

## Abstract

Neocortical computation emerges from the dynamic interplay of excitation and inhibition, operating in a loose balance regime where recurrent and external inputs contribute comparably to neuronal activity. Neurons display broad heterogeneity in synaptic inputs and firing rates, making it essential to explain the full distribution of responses, not just the mean, when elucidating mechanisms of dynamics and computation. We examined divisive normalization in mouse visual cortex using population calcium imaging of excitatory and parvalbumin (PV) inhibitory neurons, combined with computational models of varying complexity. We found that suppression in PV neurons was transiently reduced, driven by the dynamics of subcortical input, and that heterogeneity in suppression strength was linked to population correlations, variability in excitatory-inhibitory balance, and suppression of both subcortical and local cortical inputs. Our results link local recurrent connectivity to the diversity of normalizing responses in cortex, providing a mechanistic basis for functional heterogeneity in this computation.

## 1 Introduction

Neocortical computation arises from the integration of local recurrent excitation and inhibition, with each neuron receiving synapses from roughly a thousand excitatory and several hundred inhibitory cortical neurons ^1^, as well as long-range intra- and subcortical projections. These broad classes of input interact within a dynamical regime known as loose balance, in which recurrent excitation, recurrent inhibition, and external input contribute comparably on average to neuronal activity ^2^. This loose balance regime has already been shown to account for the presence of several network-level phenomena, including sublinear response summation ^3^, stimulus-evoked variability quenching ^4^, and gamma oscillations ^5^. However, functional properties exhibit substantial heterogeneity across neurons, arising in part from the heavy-tailed distributions of synaptic strengths and firing rates among their active inputs ^6;7;8^. Clarifying whether and how heterogeneity relates to loose balance could offer deeper insight into the circuit mechanisms underlying cortical computation, since a mechanistic understanding must capture both average behavior and population-wide variability.

In this work, we focus on a well-characterized sensory cortical computation: divisive normalization, which describes how neurons in sensory cortex respond sublinearly to multiple stimuli presented within their receptive fields. This computation is an appealing target for mechanistically linking circuit-level properties to the loose balance regime and functional heterogeneity to specific computational functions, as it is consistently observed across cortical areas and species ^9;10;11;12;13;14;15^. It is also supported by a range of quantitative models that provide a strong foundation for investigating underlying circuit mechanisms involving both local recurrent and external inputs ^9;16;3;17^.

The most extensively studied example of normalization is cross-orientation suppression in primary visual cortex (V1), where neurons typically exhibit sublinear summation in responses to two overlaid sine-wave gratings (i.e. plaids) ^9^. Evidence suggests that suppression in neocortex arises from a combination of altered responses in subcortical inputs and recurrent excitation and inhibition, yet the relative contributions of these components to normalization remain unresolved. Contrast saturation and rectification in the LGN, combined with spike threshold nonlinearities in V1 cells, can explain the level and variability of suppression in cortical neurons ^16;18;19;20^. Local recurrent inputs also contribute to cross-orientation suppression, as the optogenetic activation of local excitatory neurons in V1 can suppress visual responses in a manner consistent with normalization models ^21;22^. Studies of a related form of normalization, surround suppression, indicate that it is primarily mediated by the withdrawal of excitation ^23;24^ and strong inhibitory stabilization is a key feature of network models that account for excitatory withdrawal ^23;3^. Additionally, a recent network model of normalization in MT found that recurrent inhibition best explained the observed correlation patterns in that area ^25^. Together, these studies indicate a network-wide contribution to normalization.

Here, we delineate the relative contributions of these network-level mechanisms to heterogeneous normalizing responses in mouse visual cortex by imaging the dynamics of excitatory and inhibitory neurons. We interpret the results through a set of models with varying levels of circuit detail and generate new hypotheses that we then test against the data. Collectively, our findings explain how subcortical input, recurrent excitation, and recurrent inhibition jointly contribute to a heterogeneously distributed cortical canonical computation.

## 2 Results

We recorded neuronal activity in response to static sine-wave gratings at half contrast and the corresponding plaids at full contrast in the superficial layers of V1 in awake head-fixed mice using two-photon microscopy (Figure 1A, B). To image activity-dependent calcium influx, we expressed GCaMP (AAV9-Syn-GCaMP6f n=3 or -GCaMP8s n=4) panneuronally in the superficial layers of V1 in PV-Cre-Ai14 mice. The red fluorescent protein tdTomato, selectively expressed in parvalbumin-expressing (PV) interneurons through the cross of PV-Cre and Ai14 strains, allowed us to segment the imaged population into PV interneurons and non-PV, putative excitatory neurons, along with their corresponding calcium fluorescence signals. We inferred the spiking activity of each neuron from its fluorescence changes with the CASCADE algorithm ^26^, and used these inferred spike counts during stimulus presentation as our measure of the neurons’ response.

**Figure 1.**
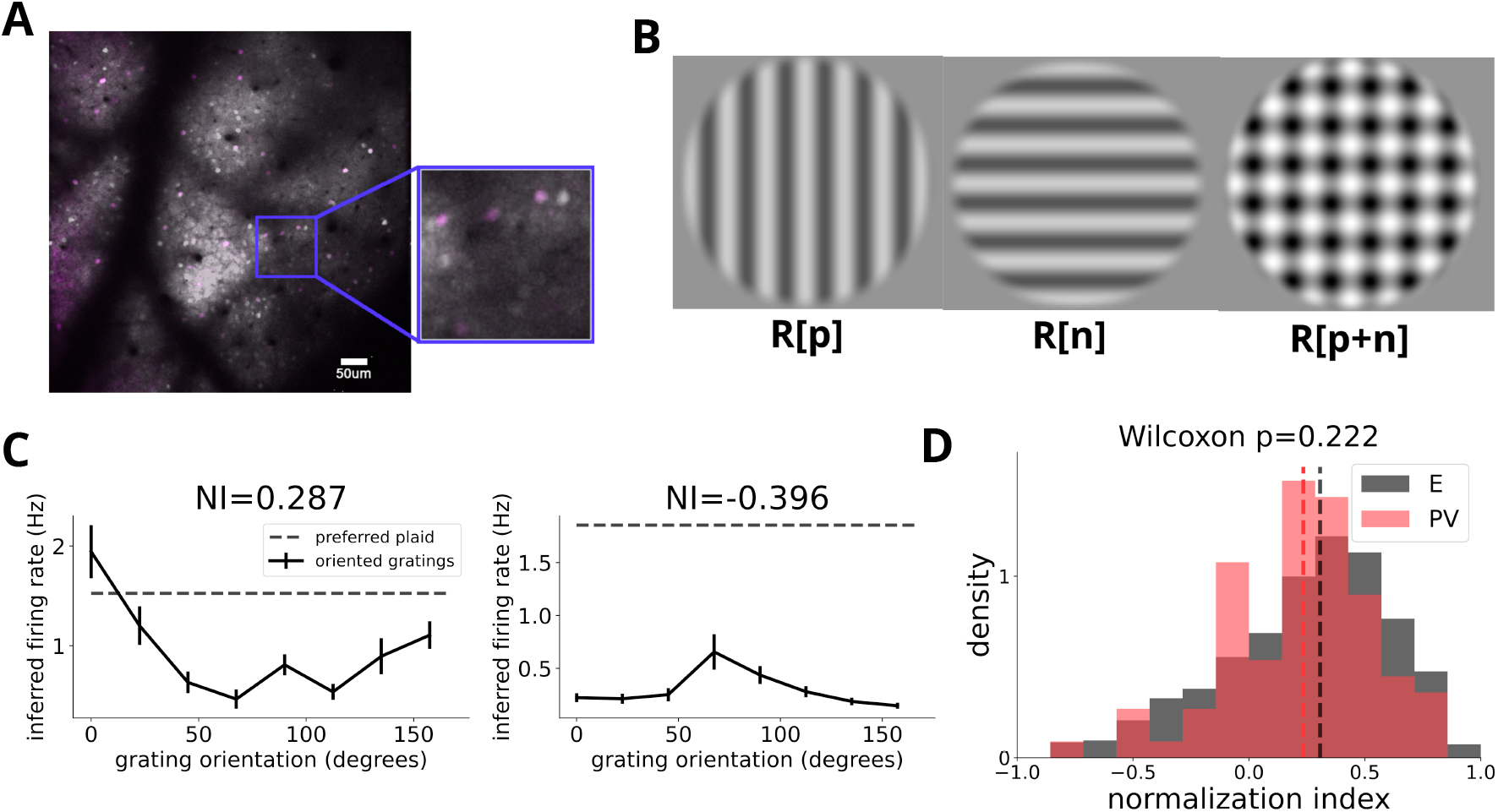
Experimental setup and basic description of normalization in mouse V1. (A) : Example imaging field of view in superficial mouse V1, with average GCAMP fluorescence in white and tdTomato expression in magenta. Colocalization of white and magenta indicates PV interneurons expressing GCaMP. (B): Example visual stimuli: large static sine-wave gratings at 50% contrast and the corresponding 100% contrast plaid resultant from summing two orthogonal gratings. (C): Example orientation tuning curves and responses to plaids at the preferred orientation for two recorded neurons, one with a positive normalization index (NI) corresponding to cross-orientation suppression, and one with a negative NI corresponding to facilitation. (D): Distribution of normalization indices (NI) for PV (N=78) and putative excitatory (N=1,325) cells pooled across mice, computed from the inferred spiking activity across the whole stimulus presentation. The median values for PV and E NI distributions, 0.23 and 0.31 respectively (indicated by dashed vertical lines), are not significantly different (*p* = 0.222, Wilcoxon rank-sum test).

We computed the degree of cross-orientation suppression in a cell’s response with the normalization index (NI) ^27^:

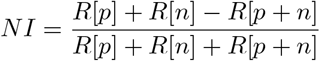

More positive NIs correspond to stronger cross-orientation suppression, and more negative NIs correspond to stronger facilitation (Figure 1C). Recurrent circuit models of normalization ^28;3^ predict that inhibitory neurons should have lower NIs on average than excitatory cells, since suppression requires that inhibition be relatively less suppressed than excitation. Both cell types exhibited a wide range of normalizing responses, consistent with previous reports ^11;20^, with moderate suppression on average (Figure 1D, median NI 0.23 for PV and 0.31 for E). However, PV and E neurons exhibited a similar range of normalization indices, with no significant difference in median from the distribution measured in excitatory cells (*p* = 0.22, Wilcoxon rank-sum test). The difference in steady-state NI between the two neuron classes may be too small to detect, or the heterogeneity in NI may be too large to allow an accurate estimate. However, because normalization is thought to shape neural responses dynamically ^17^, focusing on the dynamics of NI may prove more revealing.

### 2.1 PV interneurons show transiently lower levels of cross-orientation suppression

The inferred spikes from GCaMP8s-expressing neurons allowed sufficient temporal precision to determine the timevarying responses of cells while the slower rise time of AAV-mediated GCaMP6f fluorescence changes limited temporal precision in those mice ^29^. Therefore, we restricted our temporal response analysis to GCaMP8s-expressing mice (n=4).

We examined the time course of suppression by computing the NI separately for each 33ms imaging frame locked to onset of the one-second stimulus presentation, using the trial-averaged response in that timebin for each neuron. We observed diverse temporal response profiles across both neuronal classes—including sustained responses, onset transients, and onset-offset transients (Figure 2A). Suppression did not progress uniformly over time; instead, it was relatively lower at both stimulus onset and offset (Figure 2B). Notably, the reduction in suppression at stimulus onset was substantially greater in PV cells than in excitatory cells. During the onset transient, the median normalization index (NI) of PV cells was significantly lower than that of excitatory cells (*p* = 0.021, Wilcoxon rank-sum test, Figure 2C).

**Figure 2.**
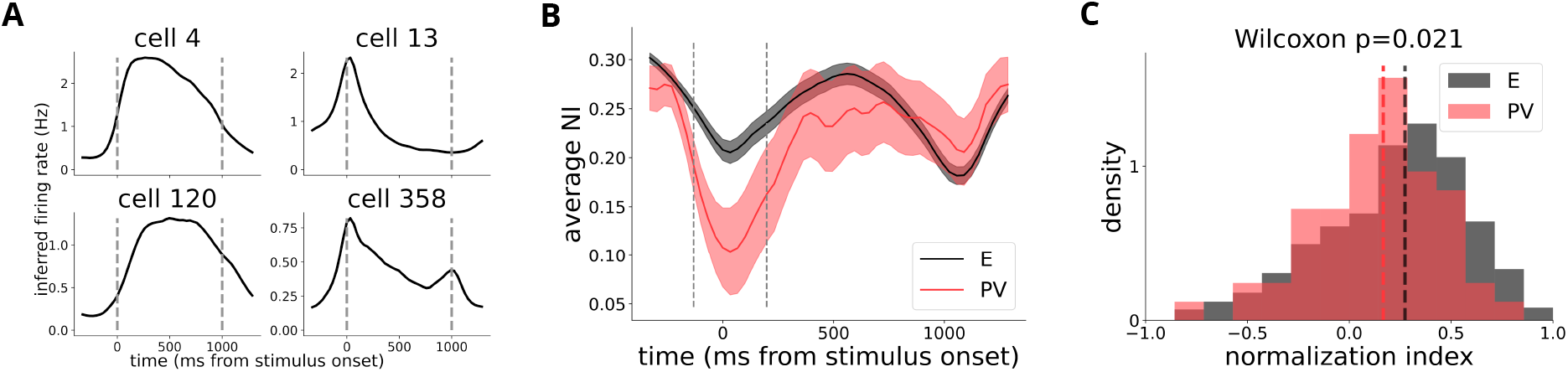
Time course of suppression reveals transiently lower suppression in PV cells. (A): Example single-unit peri-stimulus time histograms (PSTHs) constructed from inferred spiking activity across all stimuli. Vertical lines indicate stimulus onset and offset. (B): Average NI (+/-standard error) of E and PV cells as a function of time, computed from their average response at that time to their preferred and null gratings and corresponding plaid. (C)Distributions of NI computed from the summed response near stimulus onset, i.e. the area marked by vertical lines in (B) (120ms preceding through 180ms post stimulus onset), for PV (N=58) and putative excitatory (N=1,099) cells expressing GCaMP8s. This contrasts with the NI distributions in Figure 1D, which were computed from the response across the entire stimulus presentation. Medians indicated by dashed lines are 0.274 (E) and 0.167 (PV), statistically significantly lower for PV cells (*p* = 0.021, Wilcoxon rank-sum test).

Although the steady state NI of PV neurons was inconsistent with model predictions, their lower transient NI aligns with predictions from recurrent models of normalization, which predict reduced NI in inhibitory neurons ^3^. Why this effect is transient is unclear. In the related case of surround suppression, inhibitory-stabilized network models predict that suppression onset triggers a brief increase in inhibition, producing a transient (tens of milliseconds) reduction in suppression in inhibitory neurons; a prediction supported by measurements of inhibitory conductances ^23^. Alternatively, under certain parameter regimes, stabilized supralinear network (SSN) models predict a greater suppression difference between inhibitory and excitatory neurons at higher input levels ^28^.

### 2.2 Stabilized supralinear networks show transiently low NI and larger average E-I NI differences

To better understand the mechanisms underlying transiently reduced suppression in PV cells, we turned to a relatively simple model of recurrently-driven cross-orientation suppression: the ring-based stabilized supralinear network (SSN) model of normalization ^28;3^. In this network model, rate-based excitatory and inhibitory units are equally spaced along a 180-degree ring (Figure 3A), corresponding to their orientation preference. Input to the network is defined by its orientation and contrast. The network’s response properties are shaped by the expansive power-law output nonlinearities of each neuron (Figure 3B), consistent with the typical activity range of cortical neurons ^16^. Local like-to-like recurrent connectivity along the ring also contributes to network dynamics: excitatory-excitatory connections are strong enough to be unstable on their own, but stability is maintained by sufficiently strong inhibitory connectivity ^28;3^. When driven by inputs at opposite positions along the ring, corresponding to orthogonal orientations, the network units display cross-orientation suppression, since the inputs are summed sublinearly (Figure 3C).

**Figure 3.**
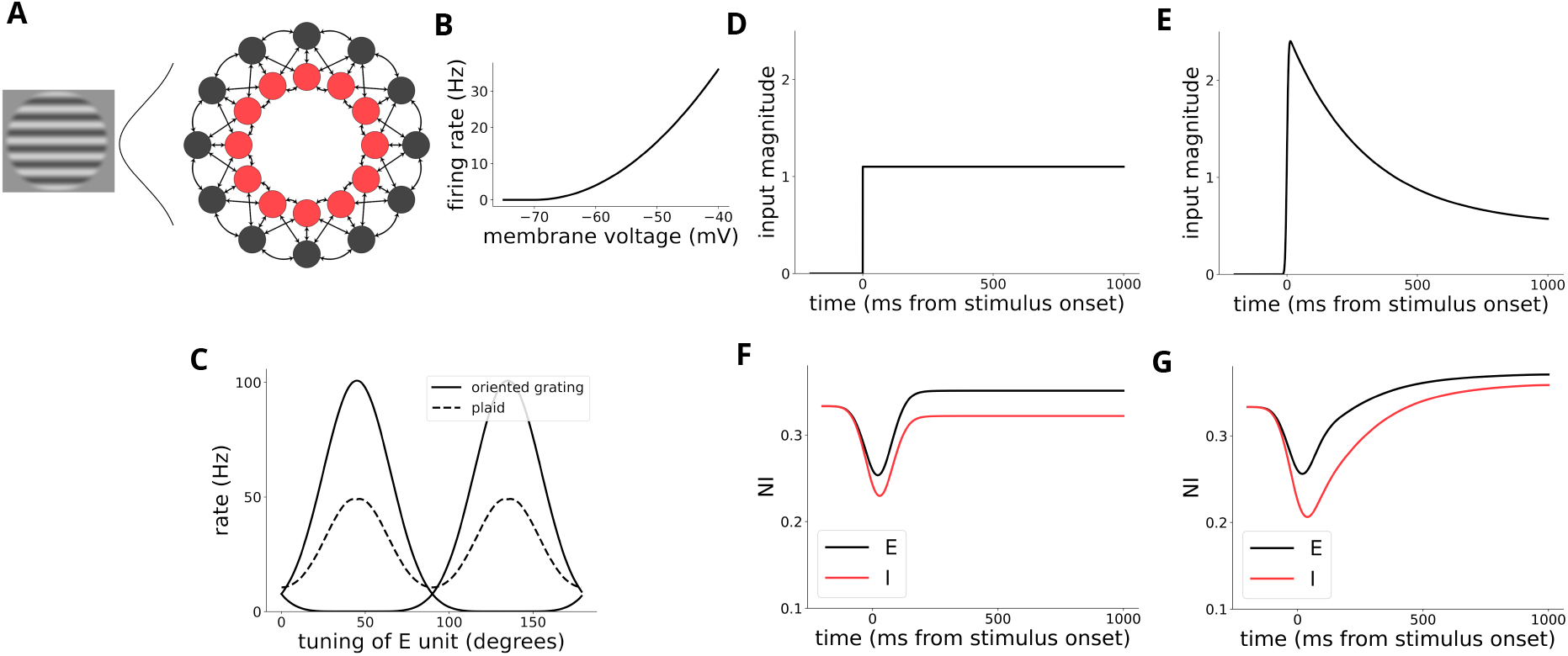
Ring SSN model shows transient NI difference with transiently high input. (A): Schematic of the ring SSN. Excitatory and inhibitory units are arrayed around the ring based on their orientation tuning. Input and recurrent connectivity take the form of Gaussian distributions around the ring. (B): The supralinear input-output nonlinearity of the units in the network. (C): Normalizing responses in the network, corresponding to 45 and 135-degree gratings, and their corresponding plaid. Units tuned to an orientation are suppressed relative to that response by the plaid. (D)and (E): Temporal input profiles for the standard flat input (D) and an exponentially decaying input (E). (F) and (G): NI over time from appropriately-tuned E and I units in the network, for the input profiles in (D) and (E). The network dynamics produce a transiently low NI as inhibition lags behind excitation even under constant input; however, when the input decays exponentially this transient is prolonged.

We examined the dynamics of normalization in the SSN model in response to two temporal profiles of input: a constant input (Figure 3D), and a transient exponentially decaying input (Figure 3E). The former isolates the dynamics of the network, while the latter qualitatively matches the time course of LGN-evoked currents in cortical cells during the presentation of static sine-wave gratings ^30^. The constant input evoked a transient reduction in the strength of suppression in both excitatory and inhibitory units, but this was brief, lasting only several tens of milliseconds, and the gap between excitatory and inhibitory NI was smaller during the transient (NI difference of 0.024) than in the steady state (0.029) (Figure 3F).The transiently decaying input produced a prolonged reduction in suppression in both cell types and amplified the difference in NI between excitatory and inhibitory units compared to the steady state (transient NI difference of 0.050 vs. 0.012 in the steady state), providing a better qualitative match to our data (Figure 3G). This suggests that the interaction between the dynamics of subcortical input and the recurrent cortical connectivity modeled by the SSN can account for the average suppression dynamics observed in V1. However, since the model is deterministic and only has one unit corresponding to each input orientation, it cannot explain the broad heterogeneity of suppression strength we observe in our data.

### 2.3 A complex V1 circuit model replicates properties of experimentally measured normalization

The ring SSN model is straightforward but fails to capture the heterogeneity in normalization strength observed in our data and does not account for the known suppression in LGN inputs to V1. ^20^. To better understand the mechanisms underlying variability in suppression, and the interactions of subcortical and recurrent mechanisms, we turned to a more complex computational model of V1: the Allen Institute’s microcircuit model ^31^. The Allen model has previously been used to probe the mechanisms behind visual flow perturbations ^32^, current source density patterns ^33^, and direction tuning ^31^. This model offers several advantages: it explicitly represents LGN input dynamics and connections, includes cortical interneuron subtypes such as PV cells, and incorporates data-driven, heterogeneous connectivity patterns with laminar detail, allowing us to focus specifically on the superficial layers for comparison with our data.

The model (Figure 4A) consists of over 200,000 model units, each a generalized leaky integrate-and-fire (GLIF) model fit to the electrophysiological properties of an individual neuronal class found in V1. Connectivity probabilities and average synaptic weights were fit to values measured through paired-patch-clamp recordings and incorporate known distance- and tuning-dependent scaling rules. The cortical model receives visual input processed by a 17,400-unit model of the LGN composed of heterogeneous spatiotemporal filters. The filters generate firing rates that are converted into Poisson spike trains and projected into the V1 model based on known thalamocortical projection patterns. The model does not include corticothalamic feedback. Further details can be found in ^31^. We stimulated the model with full-field sine-wave gratings and their corresponding plaids, to match the large stimuli presented in our *in-vivo* experiments, and analyzed the spiking of model units in response to these stimuli (Figure 4B).

**Figure 4.**
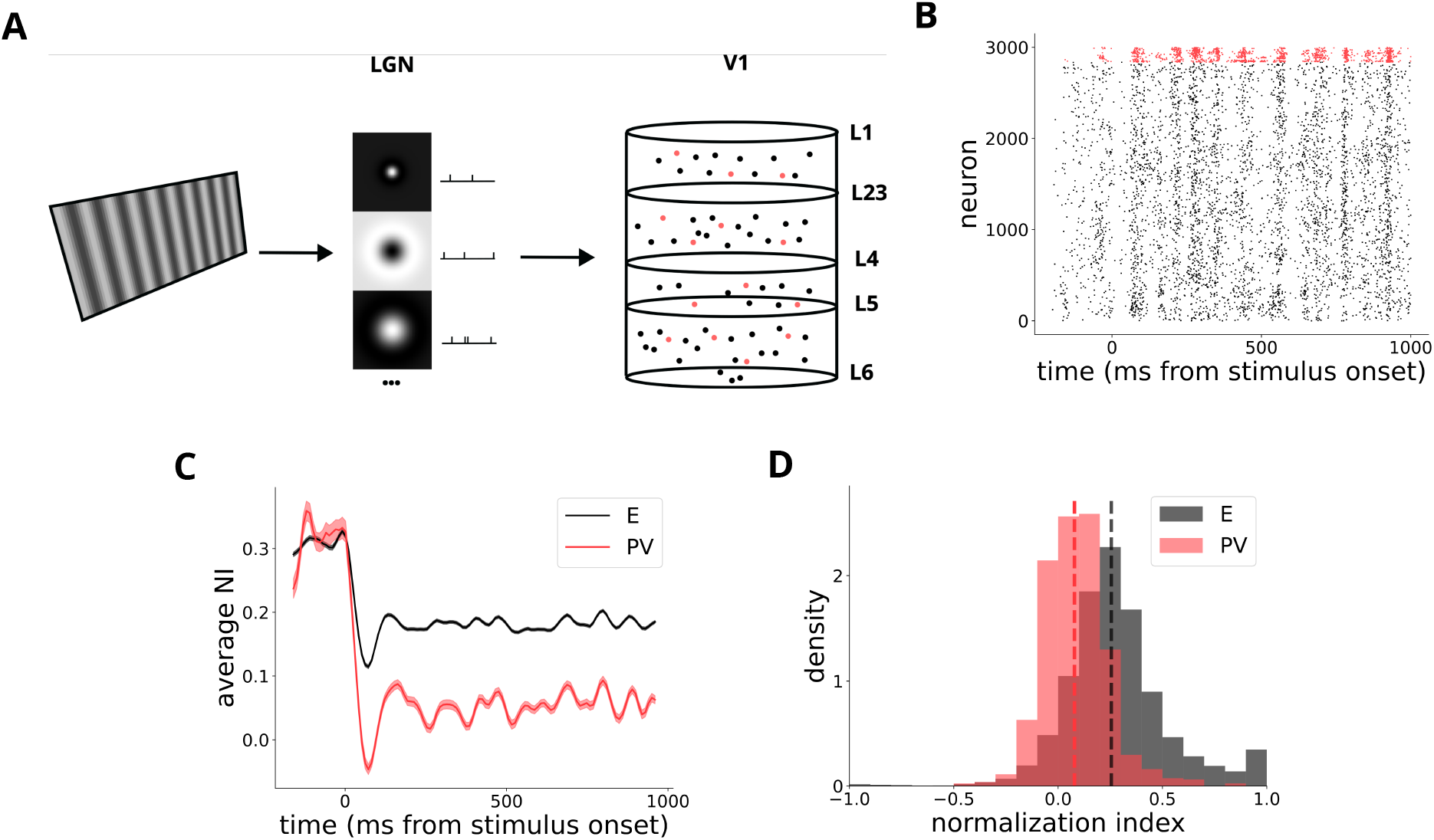
The Allen Institute V1 microcircuit model and its responses. (A): Diagram of the model. Visual input (sine-wave gratings) is fed into the LGN model, a set of spatiotemporally separable filters fit to recordings from LGN neurons. Poisson spikes generated from the firing rates output by these filters project into the V1 model, which consists of diverse cell types across all six layers of cortex. More information can be found in ^31^. (B): Raster plot of a subset of layer 2/3 excitatory and PV units (in red) from the model during a single trial. (C): Dynamics of average NI in L2/3 PV and E units in the model, transiently low at stimulus onset. Shaded error bars represent *±*1 standard error, small due to the large number of units in the model. (D): Distribution of transient NI for L2/3 excitatory and PV units in the model. Median transient NI for PV units is 0.005, much lower than the median 0.163 NI for E units (*p <* 10^−100^, Wilcoxon rank-sum test).

We measured the dynamic cross-orientation suppression in excitatory and PV units in the model’s superficial layers using the same approach as in the *in-vivo* data, calculating the normalization index (NI) from each unit’s trial-averaged responses to the preferred/null gratings and corresponding plaid. Consistent with mouse V1, the transient NI at stimulus onset in the model varied across both cell types, (Figure 4D), and the distributions of E and PV units’ transient NI largely overlapped—although the overall heterogeneity of NI in the model is lower and less skewed towards facilitation. The transient NI for PV units was lower than excitatory units’ (Figure 4D), to a greater degree than in our *in-vivo* data. The time course of normalization in both excitatory and PV model units resembled the *in-vivo* data and the ring SSN models, exhibiting a transient reduction at stimulus onset (Figure 4C). Like the SSN model, PV units in the Allen model exhibited lower levels of suppression than excitatory units throughout image presentation, but with a transiently greater reduction during the onset response, (mean ± standard error E-PV NI difference 0.151 ± 0.0064 during the transient versus 0.128 ± 0.0021 during the rest of the response.

Although not designed to study cross-orientation suppression, the Allen model captured key aspects of its heterogeneity and dynamics. We leveraged this to explore the underlying mechanisms of suppression in the model and to gain further insight into the mouse data.

### 2.4 E/I balance in the Allen model is loose, variable, and linked to suppression

Substantial evidence suggests a balance between synaptic excitation and inhibition in cortical neurons, with neurons receiving stronger excitation also receiving stronger inhibition ^34;35;36^. This balance can be “tight,” where excitation and inhibition nearly cancel, yielding a small net input, or “loose,” in which the net input is comparable in magnitude to the excitatory and inhibitory input terms ^2^. Under tight balance, average population responses scale linearly with the strength of the input, although they can appear sublinear if fully silenced neurons are considered ^37^. The SSN models considered above operate in the loose balance regime, which readily gives rise to nonlinear response properties such as cross-orientation suppression ^3;28^, because the relative level of inhibition increases with increasing network input strength. We computed a metric of E/I balance *β*, comparing each unit’s net input to its total excitatory input:

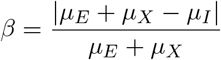

Where *µ*_*E*_ and *µ*_*I*_ denote the magnitude of intracortical excitatory and inhibitory input currents to a unit, respectively. The external input, corresponding to input from the LGN in the Allen model, is denoted by *µ*_*X*_. The balance index *β* ranges from 0 to 1, and is lower for tighter balance; *β <* 0.1 is taken as a rough cutoff for qualitatively “tight” balance ^2;38^.

The average balance in the Allen model units was overall loose, with the mean *β* = 0.86, corresponding to relatively low *µ*_*I*_ compared to *µ*_*E*_ + *µ*_*X*_. Balance was heterogeneous across units (Figure 5A), with some units receiving proportionately more or less inhibition than others, most falling between a *β* of 0.8-0.9. On a network level, SSN models generally show stronger levels of suppression under tighter balance, although the relationship depends on the precise parameters chosen ^28^. We therefore hypothesized that on a cell-by-cell level in the Allen model, heterogeneity in balance would relate to heterogeneity in suppression. Consistent with this prediction, *β* and NI were anticorrelated across units (Figure 5B, *r* = −0.15), with lower *β* (tighter balance) corresponding to stronger suppression.

**Figure 5.**
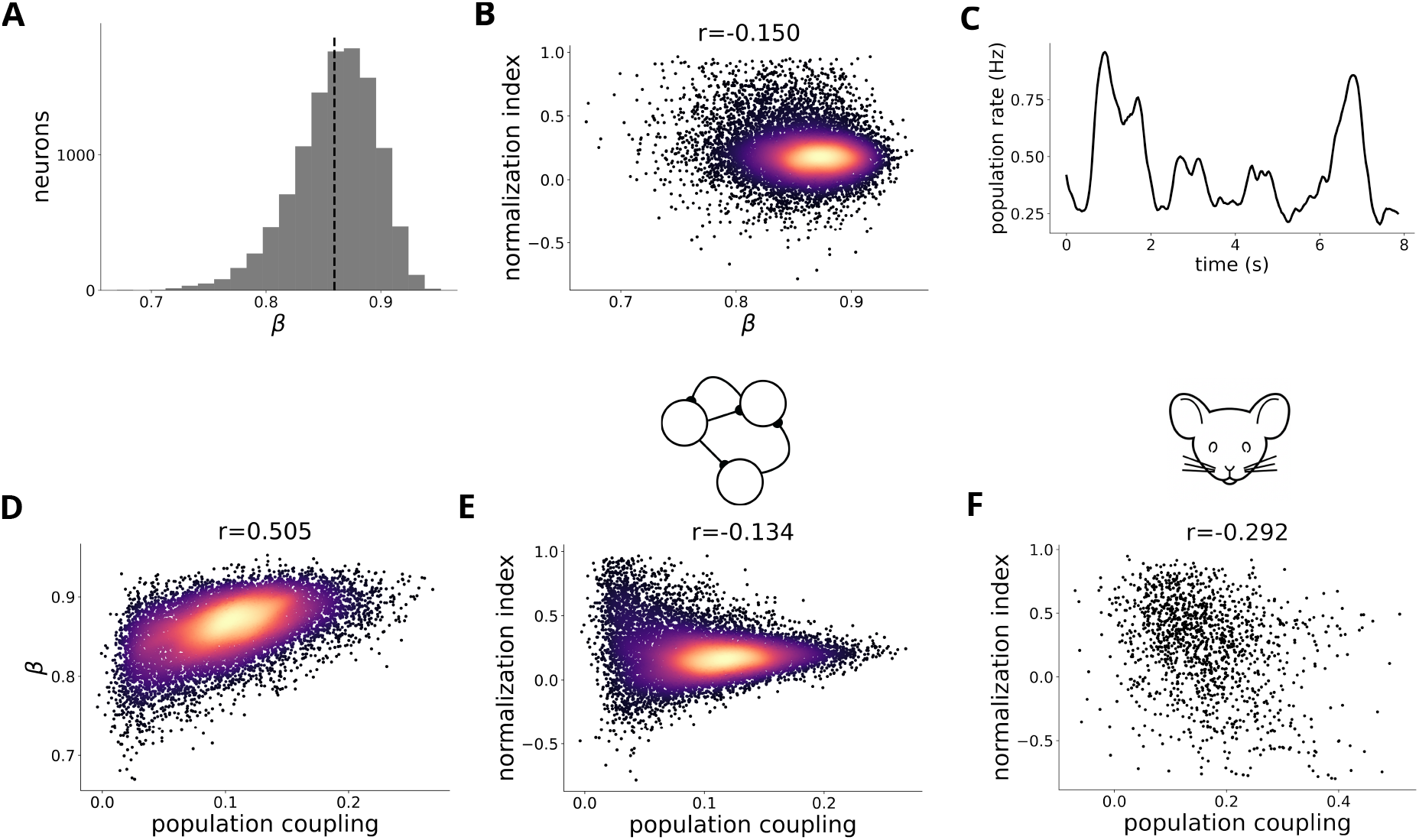
Heterogeneous E/I balance explains variability in suppression and population coupling. (A): Distribution of *β*, the measure of E/I balance, in the Allen model’s L2/3 excitatory units. (A): Suppression (measured by NI) plotted against *β, r* = −0.150, *p* = 1.3 *×* 10^−56^, corresponding to higher suppression in units with tighter E/I balance. (B): An example trace of the population rate, from the *in-vivo* data, across several seconds. Population coupling is computed as the correlation between a cell’s own time-varying activity and population rate during presentations of a given stimulus. (C): *β* plotted against population coupling in the model units, *r* = 0.505, *p <* 10^−100^. (D): NI plotted against population coupling in the model units, *r* = −0.134, *p* = 8 *×* 10^−46^. (E): NI plotted against population coupling in the *in-vivo* V1 putative excitatory cells, *r* = −0.292, *p* = 2 *×* 10^−27^.

### 2.5 E/I balance mediates a relationship between population coupling and suppression

In addition to producing nonlinear responses, loosely balanced networks more readily produce correlated variability, as tight balance results in correlations that vanish with increasing network size ^2,39;40^. Neurons that are more strongly correlated with the average activity of the local population tend to receive stronger recurrent excitatory input ^41^. Accordingly, we hypothesized that units in the model with looser E/I balance would exhibit stronger correlations with the rest of the population. Because the Allen model exhibits realistic correlated variability—typically weak, positive and tuning-dependent ^31^—we were able to test this prediction in the model units.

We measured the population coupling of cells using the correlation of their activity with the population average rate over time and across trials during stimulus presentation (Figure 5C), averaged across plaid stimuli. As predicted, we found that *β* and population coupling were positively correlated (Figure 5D, *r* = 0.51), indicating that neurons receiving relatively more excitatory input tended to be more correlated with the population. Like *β*, population coupling was weakly negatively correlated across model units with suppression (Figure 5E, *r* = −0.15), a relationship that appeared to be mediated by E/I balance—the partial correlation between coupling and NI, controlling for *β*, was substantially reduced (partial *r* = −0.07, *p* = 3.9 × 10^−13^). The relationship between *β* and population coupling was not solely explained by firing rate, despite both measures increasing with higher rates, as their partial correlation remained high after controlling for rate (partial *r* = 0.39, *p <* 10^−100^). Intuitively, units with a looser E/I balance receive more excitation from the surrounding population, making them both more coupled with population activity, and on average less suppressed by it.

These model results prompted us to test the same relationship in the *in-vivo* data, where we could compute population coupling, relate it to suppression, and thereby gain insight into the role of E/I balance in the computation of suppression. Using trial- and time-varying inferred spike traces of putative excitatory neurons from our GCaMP8s-expressing mice, we computed population coupling and found it to be negatively correlated with NI across cells, more strongly so than in the model (Figure 5F, *r* = −0.29). Although E/I balance cannot be directly measured across large populations *in-vivo*, the consistent relationship between balance and coupling simulation suggests that the observed correlation between coupling and suppression in the data may also be mediated by balance. This, in turn, points to a potential role for this recurrent network property in generating heterogeneity in cross-orientation suppression in mouse V1.

This finding is consistent with previous reports of lower pairwise correlations between more strongly normalized cells in macaque cortex ^42^, since neurons with lower population coupling tend to have lower individual pairwise correlations. A similar phenomenon was demonstrated in a recurrent circuit model of heterogeneous normalization in area MT ^25^, which found that recurrent inhibition, rather than external or local excitatory input, was the primary driver of a cell’s normalization. However, given the relative weakness of inhibition in the Allen model, such an inhibition-dominant mechanism of cross-orientation suppression seemed unlikely, suggesting that the observed relationship between normalization and coupling can arise through multiple mechanisms. To test this hypothesis, we similarly examined the mechanisms of normalization in the Allen model.

### 2.6 Feedforward and recurrent mechanisms of normalization coexist in the Allen model

We approximated the average input current to a unit from a given source—LGN, recurrent E, or recurrent I—by multiplying the trial-averaged firing rates of each of the unit’s presynaptic neighbors of that class by their synaptic strengths. By computing the trial-averaged input currents of each type for the preferred/null orientations and the corresponding plaid for a unit, we obtained the NI of each input current type. A minority of L2/3 excitatory units in the model (1,267 out of 11,099 visually responsive units) do not receive LGN input and were excluded from the analysis.

The degree of suppression of model L2/3 excitatory units was positively correlated with the degree of suppression in all three components of their inputs: subcortical, recurrent excitatory, and recurrent inhibitory (Figure 6). The strong positive correlation between NI in LGN inputs and the units’ NI mirrors the in vivo findings of ^20^ in layer 4 excitatory cells, reinforcing a subcortical contribution to suppression. A similarly strong correlation in the cortical excitatory input implies that local cortical circuitry further adds to this subcortical effect.

**Figure 6.**
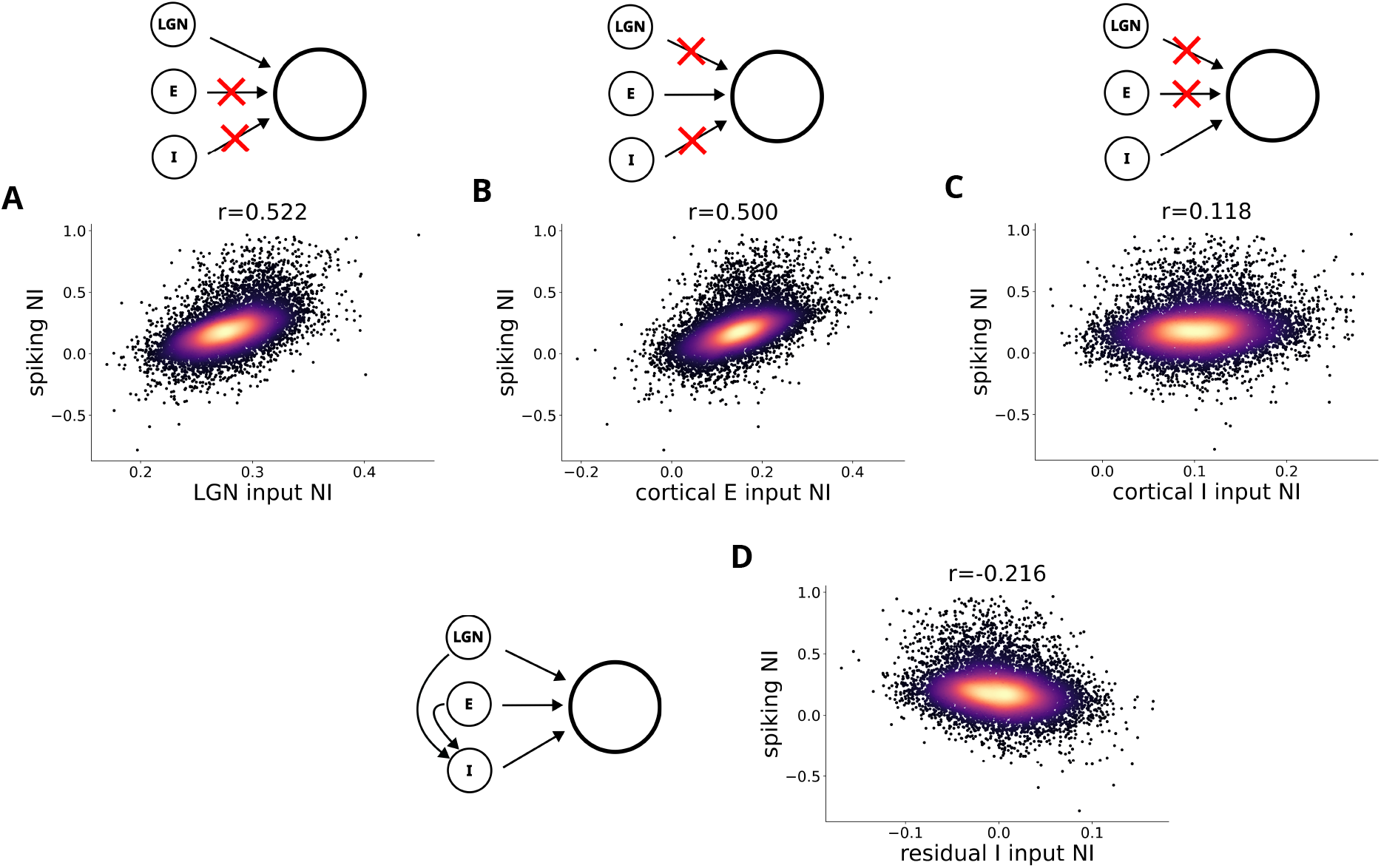
Subcortical and intracortical inputs contribute to suppression in the Allen model. (A): The suppression, measured by NI, of L2/3 E model units plotted against the suppression in their thalamic input currents, *r* = 0.522, *p <* 10^−100^. (B): Same as (A), for excitatory intracortical inputs, *r* = 0.500, *p <* 10^−100^. (C): Same as (A) and (B), for inhibitory intracortical inputs, *r* = 0.118, *p* = 8.2 *×* 10^−32^. (D): The NI of L2/3 E model units plotted against the residual of their I input NI, after regressing off both the LGN and cortical excitatory NI, *r* = −0.216, *p <* 10^−100^

The positive relationship between inhibitory input NI and spiking suppression (Figure 6C) was unexpected. All else being equal, stronger suppression of the inhibition received by a cell should lead to less suppression in the cell’s own activity, not more. Indeed, ^25^ found a strong negative correlation between the NI of a model unit’s inhibitory input and its own NI. Due to the spatial and tuning based connectivity in the Allen model, presynaptic inhibitory units tend to share excitatory inputs with the units they inhibit, potentially masking the causal effect of inhibition. To test this explanation, we computed the correlation between NI of each model unit and its inhibitory input NI after regressing off the effect of its LGN and recurrent excitatory input NI. Controlling for these shared sources of input led to a correlation of *r* = −0.218 between the residual NI of the inhibitory input and the units’ NI, recovering the expected negative relationship between inhibitory input suppression and a unit’s own suppression. However, the revealed inhibitory relationship was relatively weak compared to that of the excitatory components.

### 2.7 Recurrent excitation contributes to suppression in a heterogeneous SSN model

The strong influence of excitatory inputs on heterogeneous suppression likely reflects the weak inhibition in the Allen model, as indicated by its loose E/I balance. However, directly testing this hypothesis is challenging due to the model’s high computational demands and the difficulty of manipulating inhibitory strength in such a complex system without disrupting the stability of activity. To test whether the strength of excitation truly determines its contribution to suppression, we turned to a heterogeneous, spatially-extended SSN model adapted from ^3^, which lies between the ring SSN and the Allen model in complexity Figure 7.

**Figure 7.**
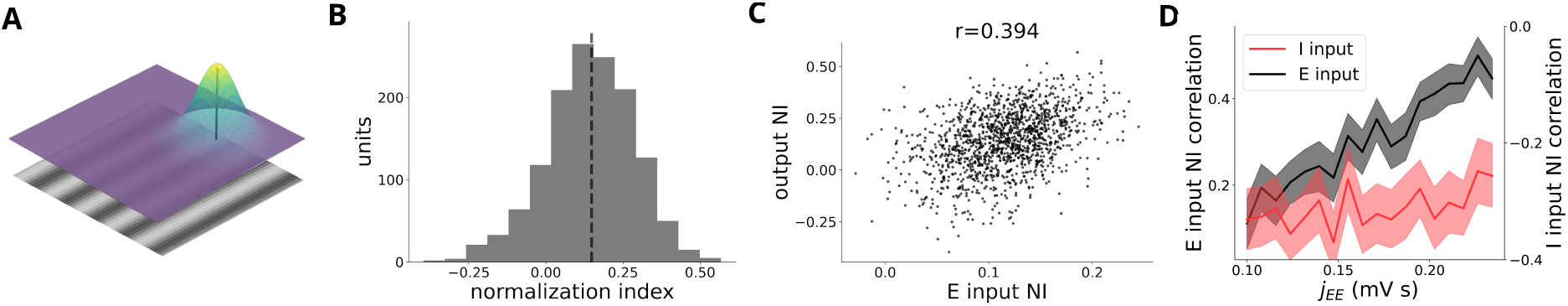
Contribution of excitatory connections to normalization in a 2D heterogeneous SSN model. (A): Schematic of the SSN’s organization in retinotopic space, with the probability density of spatial connectivity overlaid. Orientation tuning of the units at each point is randomly distributed. (B): Heterogeneous, largely positive NI in similarly-tuned units of the 2D model, qualitatively similar to the Allen model. The dashed vertical line indicates the median. (C): Example relationship between excitatory input current NI and NI in the model units, at an intermediate level of excitatory connection strength. (D): The correlation between excitatory input NI and NI across model units (left axis) increases with increasing recurrent excitatory strength, while the inhibitory input NI correlation (right axis) stays relatively flat.

Units in this model are arranged in two-dimensional retinotopic space, receive input defined by its orientation and retinotopic position, and are more strongly connected to spatially proximal and similarly orientation-tuned neighbors, as in the Allen model. While the original version of the model included spatially-organized tuning in the form of orientation columns ^3^, we randomly distributed orientation preference in space to better model the salt-and-pepper organization of mouse V1^43;44^. Instead, we measured cross-orientation suppression by presenting full-field sine-wave gratings and their corresponding plaids, matching the stimulation paradigm we used for the Allen model.

The model produces response heterogeneity because of random variation in recurrent connectivity and single-unit input–output functions. This leads to moderate variability in NI even among co-tuned units, with a distribution similar to that of the Allen model (Figure 7B). As in the Allen model, variability in a unit’s NI correlates with the NI of its inputs (Figure 7C), though only recurrent excitation and inhibition contribute, since the model’s feedforward inputs are linear by design.

We hypothesized that increasing the strength of recurrent excitatory-to-excitatory (EE) connectivity would result in a stronger correlation between the NI of their excitatory input and their own NI. This was the case: as we increased the strength of EE connectivity, the correlation between the NI of the units’ excitatory input and their own NI increased from 0.11 at the level of EE strength used in ^3^ to 0.45 at the maximum stable level of EE connectivity, while the correlation with the NI of the inhibitory input stayed flat (Figure 7D). When model units are more strongly driven by recurrent excitation, their tuning properties, specifically cross-orientation suppression, are increasingly determined by the tuning of their recurrent excitatory input. Hence, a neuron’s tuning properties are determined not only by the tuning of its inputs but also their relative strengths.

## 3 Discussion

We found that both putative excitatory and parvalbumin-expressing (PV) inhibitory neurons show transiently low levels of cross-orientation suppression at stimulus onset, rising to a higher level several hundred milliseconds after onset, and lowering again at stimulus offset. The reduction in suppression at stimulus onset was greater in PV neurons than excitatory cells. Both the onset dynamics in excitatory cells and their greater magnitude in PV cells could be explained by a transiently high, exponentially decaying input to a simple ring-based SSN model, qualitatively similar to the time course of LGN inputs to layer 4 cells during the presentation of static gratings ^30^. The nature of the offset transient was less apparent, and we did not attempt to model it mechanistically, although the lack of difference between PV and excitatory neurons at the offset transient suggest different mechanisms from the onset could be involved.

The transiently low suppression in PV interneurons, and the relationship between suppression and population coupling, support a contributing role for cortical circuitry in the computation of cross-orientation suppression in V1, rather than a purely subcortical model. Optogenetic stimulation in macaque ^21^ and ferret ^22^ further support a cortical contribution to normalizing responses. Using the Allen V1 microcircuit model, we found both subcortical and cortical contributions to cross-orientation suppression, consistent with other work supporting subcortical contributions ^45;16;46;20^. Most previous modeling work has examined either cortical or subcortical mechanisms in isolation (although see ^19^).

The models, together with corresponding in vivo measurements, revealed a central role for intracortical excitation, and particularly its relative withdrawal during plaid presentation, in generating suppression. Using a relatively simple yet heterogeneous model of cortical suppression, we found that the contribution of intracortical excitation to cross-orientation suppression increased smoothly with the strength of excitatory connections. We pursued this model and this line of inquiry because of the loose balance, driven by relatively strong excitation, we observed in the Allen model. Acknowledging that all models have limitations, the Allen model’s behavior aligned well with our in vivo measurements and generated a hypothesis that we subsequently confirmed experimentally. This may provide an explanation for the minimal effect on cross-orientation suppression of blocking intracortical inhibition reported in ^46^—as long as inhibition is sufficient to prevent runaway activity, changing levels of excitation can produce suppression. More broadly, a substantial cortical contribution to the computation may confer important advantages, including increased susceptibility to top-down modulation^47;27;48^, enabling normalization to adapt flexibly to behavioral context.

Prompted by observations in the Allen model linking heterogeneity in a cell’s balance of excitation and inhibition, its degree of correlation with the population, and its degree of suppression (Figure 5) we identified a corresponding link between the degree of suppression and population coupling in our *in-vivo* data. The tightness of E/I balance constrains the sort of computations and correlations a network can produce (as tightly balanced networks are generally linear in their inputs ^2^), and variability in this balance across cells is therefore likely to contribute to variability in the computations they perform; a possibility that has rarely been addressed (but see ^38^). The observed coupling-suppression relationship is naturally explained by the standard “normalization equation,” in which a neuron’s response is given by its input drive divided by a weighted sum of the activity of the surrounding population:

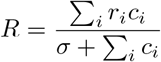

If suppression reflects the influence of the normalization denominator, which represents local population activity, then neurons showing stronger suppression should be more strongly driven by this term. Conceptually each cell can be assigned a weight *w* for the population activity term in the denominator, where a higher *w* corresponds to stronger normalization, and a negative *w* corresponds to facilitation:

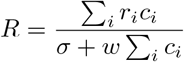

Fluctuations in population activity across time and trials would be expected to drive stronger inverse fluctuations in more strongly normalized neurons. Consequently, these neurons should fluctuate more independently of the overall population, that is, have a lower population coupling, which is precisely what we observe, further supporting the intracortical contribution to cross-orientation suppression. The relationship between the normalization equation and correlated variability, in the form of pairwise correlations, which are closely linked to population coupling ^41^, has been further explored analytically in terms of shared normalization signals in ^49;50^.

Measuring and modeling the dynamics of normalizing responses revealed transiently reduced suppression in PV interneurons and a link between suppression, excitation–inhibition balance, and population coupling, implicating local recurrent connectivity in cross-orientation suppression. By combining population imaging of excitatory and inhibitory neurons with models of varying cortical detail, we identified the relative roles of subcortical input, recurrent excitation, and recurrent inhibition. This integrated approach yields a unified framework in which these mechanisms jointly produce a canonical computation that is heterogeneously expressed across cortical populations.

## 4 Methods and Materials

Code for data processing, analysis, and simulation will be made publicly available on GitHub upon publication.

### 4.1 Mice

To selectively label PV interneurons, we crossed Cre-dependent tdTomato reporter lines (Ai14, Jax stock #007914) with PV-Cre (Jax stock #017320) C57BL/6j mice, ensuring expression of the red fluorescent protein tdTomato in PV-expressing interneurons. Animals were bred and maintained in accordance with the animal care and use regulations of the University of Chicago Institutional Animal Care and Use Committee. Male (n=5) and female (n=2) mice aged P90-P120 at the time of recording were used.

### 4.2 Surgical procedures

Mice underwent two surgeries: first viral injections, followed one or more weeks later by window and headframe implantations. For the injections, mice at least P56 were anesthetized with 1.5–2.0% isoflurane in 50% oxygen. We drilled two to three narrow holes (injection sites) in the skull centered on left-hemisphere V1 (1.5mm lateral, 0.4mm anterior of lambda). At each injection site we lowered a glass-tipped syringe controlled by a microsyringe pump into the brain and injected 200nL of AAV9 synapsin-promoted GCaMP virus (pAAV.Syn.GCaMP6f.WPRE.SV40, Addgene product #100837 for GCaMP6f mice, pGP-AAV-syn-jGCaMP8s-WPRE, Addgene product #162374 for GCaMP8s mice) at two depths, roughly 250 and 500 microns below pia. Before and after each injection, we paused for 3-5 minutes to allow the tissue to settle around the pipette and virus to diffuse more evenly.

After at least a week of recovery time, mice were again anesthetized with 1.5–2.0% isoflurane in 50% oxygen. We removed the skin on top of the animal’s skull and used dental cement to secure a custom titanium headplate to the skull. We then drilled out a 3.5mm-diameter circle of skull centered on the viral injection sites, and removed both the excised skull and the underlying dura with forceps. We then placed a 3.5mm-diameter, 0.66mm-thick glass window glued to a thin 5mm-diameter glass coverslip into the hole, so that the window was flush with the cortical surface and fully filled the space previously taken up by the skull. The wider coverslip lay on the surrounding skull and was cemented in place with dental cement.

Recordings took place 5-7 weeks after the viral injections—enough time for adequate expression of GCaMP in most neurons, but before overexpression caused a loss of dynamic signal.

### 4.3 Visual stimulation

After several days of habituation to head-fixation, we began recordings. We head-fixed mice in a custom-designed plastic sled, positioned about 23cm from a screen on their right (contralateral to the recorded area of cortex). In three mice, retinotopic mapping was performed via intrinsic signal imaging to ensure we were presenting stimuli in the appropriate location—in the rest of the mice, this was confirmed before data analysis by high levels of visual stimulus decoding.

We presented large (5̃0-degree diameter), circular, static sine-wave gratings at 50% contrast, about 0.08 cycles per degree spatial frequency, and zero phase. The corresponding plaids were the sum of two orthogonal gratings, thus at 100% contrast. All stimuli were presented for 1 second, with 1 second luminance-matched grey screens presented between. For the three GCaMP6f mice, we presented 4 evenly-spaced grating stimuli and their 2 corresponding plaids for a total of 175 trials each. For the four GCaMP8s mice, we presented 8 evenly-spaced grating stimuli and their 4 corresponding plaids for a total of 104 trials each. The stimuli were presented in 8 imaging blocks of 5 minutes each, in random interleaved order within each block, with one minute of luminance-matched grey between blocks. Stimulus presentation was handled with custom Python scripts utilizing the PsychoPy package ^51^.

### 4.4 Data collection and analysis

We performed calcium imaging with a custom-built two-photon microscope, controlled with the ScanImage Basic software ^52^. A 16x Olympus objective provided a field of view (FOV) of roughly 750×750 microns.

We used the Suite2p software ^53^ to extract ROIs from the raw imaging video, which were then classified as cells or non-cell ROIs based on a classifier trained on several manually-curated FOVs. We subtracted the signal from the surrounding neuropil with a varying coefficient, estimated for each cell based on optimizing the smoothness of the resulting trace, modified from the Allen Institute’s pipeline code at https://github.com/AllenInstitute/ophys_etl_pipelines/tree/main. Then, we computed traces of dF/F, the percentage change from baseline fluorescence, for each cell, estimating the baseline with a median filter-based approach modified from the Allen Institute’s pipeline. Finally, these dF/F traces were used as input for the CASCADE algorithm ^26^, using the “GC8s EXC 30Hz smoothing50ms high noise” models for the 8s recordings and the “Global EXC 30Hz smoothing50ms high noise” models for the 6f recordings, resulting in inferred spike counts across the recordings for each neuron. The CASCADE algorithm is trained on smoothed ground-truth spikes (in our case, 50ms-width-Gaussian smoothed), and “looks ahead” in time for optimal performance; hence its inferred spike traces can sometimes precede the actual spikes, though on average, they should not. This explains why the rise in the PSTHs in Figure 2A and the transient drop in NI in Figure 2C precede the stimulus onset; even though their peak values always occur post-onset.

For the analyses of visual responses, we only included neurons that passed a test of visual responsiveness. This required that they respond at least 10% more strongly on average during stimulus presentation, averaged across all stimuli, compared to the between-stimulus greys, and additionally that this stimulus-evoked response was statistically significantly higher than the between-stimulus response at a *p* = 0.05 level from a one-sided t-test. Statistical tests were computed using the associated functions in the SciPy package ^54^, and general analysis heavily used NumPy ^55^.

For the full-response normalization indices displayed in Figure 1, we summed the inferred spike count across all 30 imaging frames corresponding to a single second-long presentation, and used this number as the response for a cell to that stimulus trial. NIs were then computed from trial-averaged responses to each of the corresponding stimuli for a cell, picking the preferred grating as the orientation that evoked the maximal trial-averaged response in the cell.

Population coupling was computed from the time-varying traces during all plaid stimuli. For each plaid stimulus presentation, we extracted the 30 imaging frames corresponding to its presentation, and concatenated these end-to-end, to capture the time- and trial-varying responses. We then computed the population average trace for each cell by averaging across the rest of the population, excluding that cell, and computed the population coupling from the correlation of the cell’s trace with that population trace across all time and trials. Application of this method to electrophysiological data introduces a rate bias ^41^, with higher firing-rate cells showing higher coupling. We observed a similar correlation with rate in our data, but found that the proposed alternative, the “population spike-triggered average” method, was also rate-biased. Therefore, we use the simpler correlation-based method.

For the temporal analyses of NI in Figure 2, we compute the average inferred spike count at each time bin for the relevant stimuli when computing NI for each cell, with the preferred orientation selected from its time-averaged response. We then compute the NI for each cell independently from the trial-averaged response in each onset-locked imaging frame, and report the average NI across cells of a particular type during that frame. The transient NIs in Figure 2 C are computed from the average response across the time bins corresponding to the transient (from 120ms pre-stimulus onset to 180ms post-onset), rather than averaging the NI computed from each time bin separately.

### 4.5 SSN ring model

The ring SSN model was adapted from ^3^ and ^4^, not including the stochastic variant in the latter. The dynamics described here largely adapted the terms from ^4^. Units were positioned, with even spacing, around the 180-degree ring, with an excitatory and inhibitory neuron at each position, for a total of *N* = *N*_*E*_ + *N*_*I*_ units. Units followed the dynamics:

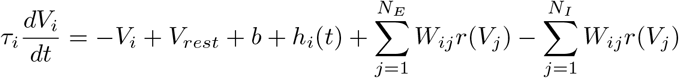

Where *V*_*i*_ indicates the membrane voltage of unit *i, τ*_*i*_ its time constant, *h*_*i*_(*t*) its feedforward input, and *W*_*ij*_ is the connection weight from unit *j* to unit *i*. The baseline stimulus-independent input *b* was shared across all units. See Table 1 for parameter values.

**Table 1.**
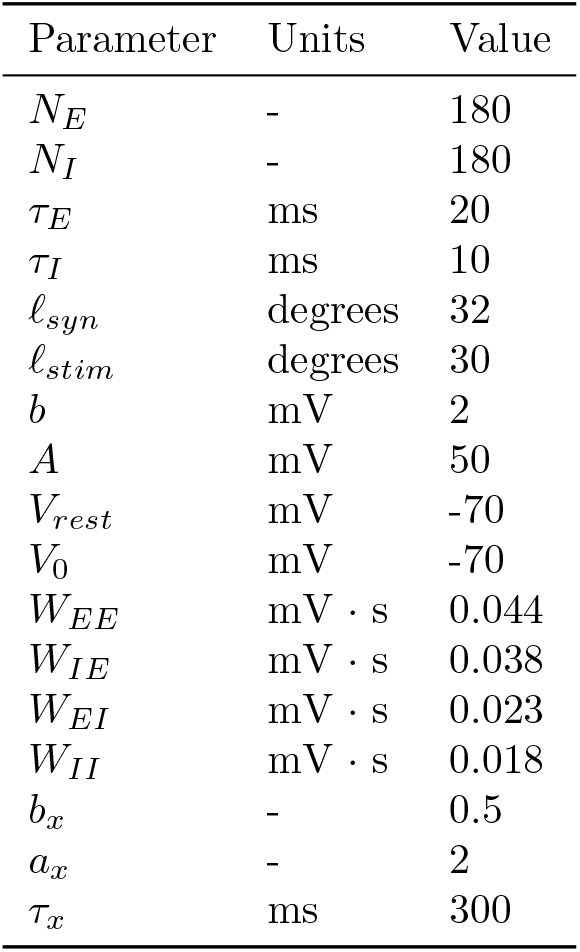
Parameter values for the ring SSN.

| Parameter | Units | Value |
| --- | --- | --- |
| $N_E$ | - | 180 |
| $N_I$ | - | 180 |
| $\tau_E$ | ms | 20 |
| $\tau_I$ | ms | 10 |
| $\ell_{syn}$ | degrees | 32 |
| $\ell_{stim}$ | degrees | 30 |
| $b$ | mV | 2 |
| $A$ | mV | 50 |
| $V_{rest}$ | mV | -70 |
| $V_0$ | mV | -70 |
| $W_{EE}$ | mV · s | 0.044 |
| $W_{IE}$ | mV · s | 0.038 |
| $W_{EI}$ | mV · s | 0.023 |
| $W_{II}$ | mV · s | 0.018 |
| $b_x$ | - | 0.5 |
| $a_x$ | - | 2 |
| $\tau_x$ | ms | 300 |

The symmetric weights *W*_*ij*_ between units *i* and *j* of classes *α* and *β* were:

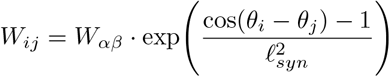

Where *α, β* ∈ {*E, I*}.

The supralinear input-output function of the units, *r*, was defined as the rectified power-law of its membrane voltage:

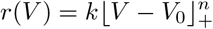

Where *V* is the membrane voltage, and the gain *k*, exponent *n*, and threshold *V*_0_ are given in Table 1.

The time-varying input to the *i*th network unit during the presentation of a given orientation *θ*_*stim*_ was given as:

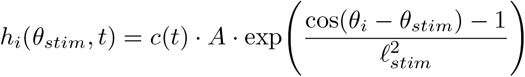

Where *A* is the scale of the input, *ℓ*_*stim*_ is the length constant of the input bump, and the time-varying input strength *c*(*t*) took different forms based on the input form. For the constant input, *c*(*t*) = 0 for *t <* 0, and *c*(*t*) = 1.1 for *t* ≥ 0, with the value of 1.1 chosen to roughly match the average value of the time-varying input. For the time varying input, we have, for *t >* 0:

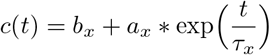

Where *b*_*x*_, *a*_*x*_, and *τ*_*x*_ are the baseline, maximum amplitude, and time constant of the exponentially decaying input. The *c*(*t*) trace is additionally smoothed with a 5ms-width Gaussian filter to remove artifacts in the network response from the sharp jump up at *t* = 0.

For each simulation, we initialize the model at *V*_*m*_ = *V*_*rest*_ for all units, and run it for a 200ms “warmup” period to let it reach a steady state with no input beyond the baseline input *b*. Then we present it with the input traces *h*(*t*) shown in Figure 3D and E. We compute the NI from the responses of neurons tuned to 45^°^ when presented with stimuli at 45^°^ (the preferred orientation), 135^°^ (the null orientation), and the sum of those two inputs (the plaid). The dynamic NI curves presented in Figure 3F and G are additionally smoothed by a 50ms-width Gaussian to mimic the smoothing process implicit in our *in-vivo* data.

### 4.6 Allen Institute model

A detailed description of the organization of the Allen Institute V1 microcircuit model can be found in ^31^. Here, we describe only our use of the GLIF-based version of the model, and the analyses we performed on the responses of its units. We used the optimized weights for the GLIF model downloaded from the Dropbox folder provided by the Allen Institute at https://portal.brain-map.org/explore/models/mv1-all-layers, and ran it using the BMTK software from https://github.com/AllenInstitute/bmtk^56^, compiled locally with MPI support. Running the model on a 32-core server with 252 GB of RAM, with 8 parallel processes, simulating 1.5 seconds of activity (the length of our trials) took roughly two minutes of wall time.

#### 4.6.1 Visual stimulation for the model

We stimulated the model with full-field drifting gratings at 50% contrast and their corresponding plaids at 100% contrast, evenly spaced at 16 different directions from 0 to 360 degrees, drifting at a temporal frequency of 2 Hz. While we used static gratings in our *in-vivo* experiments, we found that the model did not respond well to static gratings, possibly reflecting the fact that it was optimized to better fit experimental values of direction selectivity than pure orientation selectivity. We chose to always use the 90-degree-clockwise rotation of a grating as the “null” grating to construct the plaid, so we had a total of 32 stimuli presented to the model: 16 gratings at different orientations, and the 16 plaids corresponding to each grating overlaid with its 90-degree-clockwise-rotated null.

For each stimulus, we simulated 100 independent trials from the LGN model, which generates variable spike trains from an inhomogenous Poisson process for each LGN cell. Trials lasted for 1,500 ms of model time, presenting a luminance-matched grey for the first 500 ms and presenting the appropriate stimulus for the last second. Stimuli were presented at the same phase for each trial, except for the data in Figure 4D. For that analysis we found a phase-locked temporal modulation of the average NI matching the drifting frequency, so we randomized the phase trial-by-trial to eliminate it. The V1 model’s responses for each trial were simulated with that trial’s LGN spikes as input, as well as an independent realization of the 1 kHz shared background signal.

#### 4.6.2 Analysis of the model’s responses

Our analyses of the model units focused on the excitatory and PV units in the superficial layers of the model, which are treated as a single entity, “Layer 2/3”. In this layer, there were 12,689 excitatory and 640 PV units within the center 400 *µ*m of the model. We restricted our analysis to units that passed the same measure of visual responsiveness used for the *in-vivo* data—a minimum 10% and statistically significant increase in response to visual stimuli over the luminance-matched grey. For analyses relating to normalization indices, we excluded the small minority of excitatory units that did not spike at all in response to one of the stimuli used to compute the NI (resulting in an NI of −1 or 1 exactly). After these two steps of sub-selection, we were left with 11,099 excitatory units.

For analyses of temporally-varying responses in Figure 4 and population coupling in Figure 5, we binned spikes at 10 ms and smoothed the resulting traces with a 15 ms-width Gaussian filter. The transient response in Figure 4C was defined as the time period from 70ms post-stimulus onset to 120ms post-onset, based on visual observation of the time course of response in the model. The “sustained” portion of the response was defined as 120 ms post-onset to the end of presentation. Population coupling was computed from 40 ms post-stimulus onset to the end of presentation, separately for each stimulus, and the correlations for each neuron were averaged across all the plaid stimuli, consistent with analysis of neural data.

#### 4.6.3 Analysis of model inputs

For the computation of *β* in Figure 5 as well as the NI of various input types in Figure 6, we approximated the average input current to a unit from a given source, during stimulus, as the average spike count of the source multiplied by its weight to the unit. For example, the computation of *µ*_*E*_ for some unit *j* with *N*_*p*_ cortical pre-synaptic excitatory units, during the presentation of stimulus *s*, would be

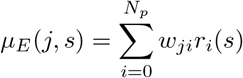

Where *w*_*ji*_ is the strength of the weight in the model between unit *j* and presynaptic unit *i*, and *r*_*i*_ is the trial-averaged spike count of unit *i* in response to stimulus *s*. For the computation of *β, µ*_*I*_ was additionally multiplied by 1.47 to account for the slower dynamics of inhibitory synapses in the model (a given weight for an inhibitory synapse results in about 1.47 times the input current as an excitatory synapse with the same weight). This method is similar to the “effective connectivity” used to analyze the Allen model’s mechanisms of visual flow responses in ^32^.

The normalization index of each input source (LGN, cortical excitation, and cortical inhibition) was computed from the total input current for that source in response to the preferred grating (and corresponding null/plaid) for the post-synaptic neuron.

### 4.7 Heterogeneous SSN model

The heterogeneous salt-and-pepper SSN was adapted from the probabilistically connected 2D SSN model in ^3^. Model parameters can be found in Table 2. Units followed the same dynamics and supralinear input-output function as the ring SSN.

**Table 2.** Parameter values for the heterogeneous SSN.

| Parameter | Units | Value |
| --- | --- | --- |
| $N_E$ | - | 75 |
| $N_I$ | - | 75 |
| $\tau_E$ | ms | 20 |
| $\tau_I$ | ms | 10 |
| $\sigma_o$ | spatial degrees | 45 |
| $\Delta x$ | spatial degrees | $\frac{16}{N_E}$ |
| $\kappa_E$ | - | 0.1 |
| $\kappa_I$ | - | 0.5 |
| $\sigma_{EE}$ | tuning degrees | $8\Delta x$ |
| $\sigma_{EI}$ | tuning degrees | $4\Delta x$ |
| $\sigma_{IE}$ | tuning degrees | $12\Delta x$ |
| $\sigma_{II}$ | tuning degrees | $4\Delta x$ |
| $\ell_{stim}$ | tuning degrees | 32 |
| $J_{EE}$ | mV · s | 0.1 to 0.23 |
| $J_{IE}$ | mV · s | 0.18 |
| $J_{EI}$ | mV · s | 0.089 |
| $J_{II}$ | mV · s | 0.096 |
| $b$ | mV | 0 |
| $A$ | mV | 25 |
| $V_{rest}$ | mV | 0 |
| $V_0$ | mV | 0 |
| $n$ | - | 2.0 |
| $k$ | - | 0.012 |

The major difference between the heterogeneous salt-and-pepper SSN and the ring SSN is the spatial structure of the former. Excitatory and inhibitory units are laid out in an *N*_*E*_ × *N*_*E*_ grid evenly spaced across 16 × 16 degrees of visual space, with *N*_*E*_ the number of excitatory neurons and the number of inhibitory neurons *N*_*I*_ = *N*_*E*_. We constructed the orientation tuning following the methods of ^3^, using the algorithm for orientation column generation from ^57^, but then randomized the resulting values in space, to simultaneously match the overall statistics of orientation tuning and also match the unorganized, “salt-and-pepper” arrangement of orientation tuning in mouse V1. This meant that units with preferred orientation *θ* were randomly distributed retinotopically.

Nonzero recurrent connections in the network were sparse, and randomly drawn, for a presynaptic neuron *j* and postsynaptic unit *i* of classes *β* and *α* ∈ {*E, I*} respectively, according to:

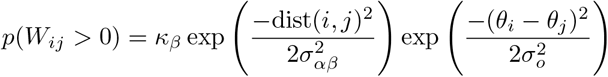

Where *σ*_*αβ*_ is the spatial width of connectivity from class *β* to *α, σ*_*o*_ is the shared width of orientation-tuned connectivity, and the spatial distance function “dist” is Euclidean distance in degrees, with periodic conditions at the boundaries of the grid. Drawn connections were then randomly assigned weights *W*_*ij*_ ~ 𝒩 (*J*_*αβ*_, (0.25*J*_*αβ*_)^2^), which were then normalized by the average number of connections so that the total incoming weight to neuron *i* was *J*_*αβ*_.

This process produced heterogeneity in recurrent connectivity, but unit-by-unit heterogeneity was additionally provided by drawing the parameters *τ*, and *n* for each neuron from normal distributions with a standard deviation 0.05 their mean.

The network was simulated for 250 ms with timesteps of 0.1 ms, by which point it settled to a steady state. We analyzed responses from the last timestep of simulation in response to full-field gratings at 0 and 90 degrees, and their corresponding plaid. We restricted the analysis of the relationship between inputs and outputs to excitatory units tuned to within 10 degrees of either 0 or 90 degrees.

## 5 Acknowledgments

This work was supported by the BRAIN Initiative’s Targeted Brain Circuit Projects under award RO1 121772. HR was additionally supported by NIH training grant 5R90DA060338-02/5T90DA059109-02 Training in Theory and Computation for Next Generation Neuroscientists. We would like to thank John Maunsell, Chery Cherian, Gabriella Wheeler Fox and Maria Pope for helpful discussions and feedback on the manuscript.

## References

[1] Gregg Wildenberg, Hanyu Li, Vandana Sampathkumar, Anastasia Sorokina, and Narayanan Kasthuri. Isochronic development of cortical synapses in primates and mice. Nature Communications, 14(1):8018, 2023.

[2] Yashar Ahmadian and Kenneth D. Miller. What is the dynamical regime of cerebral cortex? Neuron, 109(21):3373–3391, 2021. ISSN 0896-6273. doi: 10.1016/j.neuron.2021.07.031. URL https://www.sciencedirect.com/science/article/pii/S0896627321005754.

[3] Daniel B Rubin, Stephen D Van Hooser, and Kenneth D Miller. The stabilized supralinear network: a unifying circuit motif underlying multi-input integration in sensory cortex. Neuron, 85(2):402–417, 2015.

[4] Guillaume Hennequin, Yashar Ahmadian, Daniel B Rubin, Máté Lengyel and Kenneth D Miller. The dynamical regime of sensory cortex: stable dynamics around a single stimulus-tuned attractor account for patterns of noise variability. Neuron, 98(4):846–860, 2018.

[5] Rodrigo Echeveste, Laurence Aitchison, Guillaume Hennequin, and Máté Lengyel. Cortical-like dynamics in recurrent circuits optimized for sampling-based probabilistic inference. Nature neuroscience, 23(9):1138–1149, 2020.

[6] Sen Song, Per Jesper Sjöström, Markus Reigl, Sacha Nelson, and Dmitri B Chklovskii. Highly nonrandom features of synaptic connectivity in local cortical circuits. PLoS biology, 3(3):e68, 2005.

[7] György Buzsáki and Kenji Mizuseki. The log-dynamic brain: how skewed distributions affect network operations. Nature Reviews Neuroscience, 15(4):264–278, 2014.

[8] Tomáš Hromádka, Michael R DeWeese, and Anthony M Zador. Sparse representation of sounds in the unanesthetized auditory cortex. PLoS biology, 6(1):e16, 2008.

[9] David J Heeger. Normalization of cell responses in cat striate cortex. Visual neuroscience, 9(2):181–197, 1992.

[10] Matteo Carandini, David J Heeger, and J Anthony Movshon. Linearity and normalization in simple cells of the macaque primary visual cortex. Journal of Neuroscience, 17(21):8621–8644, 1997.

[11] Zaina A Zayyad, John HR Maunsell, and Jason N MacLean. Normalization in mouse primary visual cortex. Plos one, 18(12):e0295140, 2023.

[12] Davide Zoccolan, David D Cox, and James J DiCarlo. Multiple object response normalization in monkey inferotemporal cortex. Journal of Neuroscience, 25(36):8150–8164, 2005.

[13] Eero P Simoncelli and David J Heeger. A model of neuronal responses in visual area mt. Vision research, 38 (5):743–761, 1998.

[14] Neil C Rabinowitz, Ben DB Willmore, Jan WH Schnupp, and Andrew J King. Contrast gain control in auditory cortex. Neuron, 70(6):1178–1191, 2011.

[15] Gijs Joost Brouwer, Vanessa Arnedo, Shani Offen, David J Heeger, and Arthur C Grant. Normalization in human somatosensory cortex. Journal of Neurophysiology, 114(5):2588–2599, 2015.

[16] Nicholas J Priebe and David Ferster. Mechanisms underlying cross-orientation suppression in cat visual cortex. Nature neuroscience, 9(4):552–561, 2006.

[17] David J Heeger and Klavdia O Zemlianova. A recurrent circuit implements normalization, simulating the dynamics of v1 activity. Proceedings of the National Academy of Sciences, 117(36):22494–22505, 2020.

[18] Baowang Li, Jeffrey K Thompson, Thang Duong, Matthew R Peterson, and Ralph D Freeman. Origins of cross-orientation suppression in the visual cortex. Journal of Neurophysiology, 96(4):1755–1764, 2006.

[19] Melinda Koelling, Robert Shapley, and Michael Shelley. Retinal and cortical nonlinearities combine to produce masking in v1 responses to plaids. Journal of computational neuroscience, 25:390–400, 2008.

[20] Dylan Barbera, Nicholas J. Priebe, and Lindsey L. Glickfeld. Feedforward mechanisms of cross-orientation interactions in mouse v1. Neuron, 110(2):297–311.e4, 2022. ISSN 0896-6273. doi: 10.1016/j.neuron.2021.10.017. URL https://www.sciencedirect.com/science/article/pii/S0896627321007856.

[21] Jonathan J Nassi, Michael C Avery, Ali H Cetin, Anna W Roe, and John H Reynolds. Optogenetic activation of normalization in alert macaque visual cortex. Neuron, 86(6):1504–1517, 2015.

[22] Shen Wang, Agostina Palmigiano, Kenneth D Miller, and Stephen D Van Hooser. Targeted cortical stimulation reveals principles of cortical contextual interactions. bioRxiv, pages 2022–06, 2022.

[23] Hirofumi Ozeki, Ian M Finn, Evan S Schaffer, Kenneth D Miller, and David Ferster. Inhibitory stabilization of the cortical network underlies visual surround suppression. Neuron, 62(4):578–592, 2009.

[24] Tatsuo K Sato, Bilal Haider, Michael Häusser, and Matteo Carandini. An excitatory basis for divisive normalization in visual cortex. Nature neuroscience, 19(4):568–570, 2016.

[25] Deying Song, Douglas Ruff, Marlene Cohen, and Chengcheng Huang. Neuronal heterogeneity of normalization strength in a circuit model. bioRxiv, pages 2024–11, 2024.

[26] Peter Rupprecht, Stefano Carta, Adrian Hoffmann, Mayumi Echizen, Antonin Blot, Alex C. Kwan, Yang Dan, Sonja B. Hofer, Kazuo Kitamura, Fritjof Helmchen, and Rainer W. Friedrich. A database and deep learning toolbox for noise-optimized, generalized spike inference from calcium imaging. Nature Neuroscience, 24(9):1324–1337, Sep 2021. ISSN 1546-1726. doi: 10.1038/s41593-021-00895-5. URL https://doi.org/10.1038/s41593-021-00895-5.

[27] Amy M Ni, Supratim Ray, and John HR Maunsell. Tuned normalization explains the size of attention modulations. Neuron, 73(4):803–813, 2012.

[28] Yashar Ahmadian, Daniel B Rubin, and Kenneth D Miller. Analysis of the stabilized supralinear network. Neural computation, 25(8):1994–2037, 2013.

[29] Peter Rupprecht, Márton Rózsa, Xusheng Fang, Karel Svoboda, and Fritjof Helmchen. Spike inference from calcium imaging data acquired with gcamp8 indicators. bioRxiv, pages 2025–03, 2025.

[30] Anthony D Lien and Massimo Scanziani. Cortical direction selectivity emerges at convergence of thalamic synapses. Nature, 558(7708):80–86, 2018.

[31] Yazan N Billeh, Binghuang Cai, Sergey L Gratiy, Kael Dai, Ramakrishnan Iyer, Nathan W Gouwens, Reza Abbasi-Asl, Xiaoxuan Jia, Joshua H Siegle, Shawn R Olsen, et al. Systematic integration of structural and functional data into multi-scale models of mouse primary visual cortex. Neuron, 106(3):388–403, 2020.

[32] Javier Galván Fraile, Franz Scherr, José J Ramasco, Anton Arkhipov, Wolfgang Maass, and Claudio R Mirasso. Modeling circuit mechanisms of opposing cortical responses to visual flow perturbations. PLOS Computational Biology, 20(3):e1011921, 2024.

[33] Atle E Rimehaug, Alexander J Stasik, Espen Hagen, Yazan N Billeh, Josh H Siegle, Kael Dai, Shawn R Olsen, Christof Koch, Gaute T Einevoll, and Anton Arkhipov. Uncovering circuit mechanisms of current sinks and sources with biophysical simulations of primary visual cortex. elife, 12:e87169, 2023.

[34] Yousheng Shu, Andrea Hasenstaub, and David A McCormick. Turning on and off recurrent balanced cortical activity. Nature, 423(6937):288–293, 2003.

[35] Michael Okun and Ilan Lampl. Instantaneous correlation of excitation and inhibition during ongoing and sensory-evoked activities. Nature neuroscience, 11(5):535–537, 2008.

[36] Mingshan Xue, Bassam V Atallah, and Massimo Scanziani. Equalizing excitation–inhibition ratios across visual cortical neurons. Nature, 511(7511):596–600, 2014.

[37] Cody Baker, Vicky Zhu, and Robert Rosenbaum. Nonlinear stimulus representations in neural circuits with approximate excitatory-inhibitory balance. PLoS computational biology, 16(9):e1008192, 2020.

[38] Jonathan F O’Rawe, Zhishang Zhou, Anna J Li, Paul K LaFosse, Hannah C Goldbach, and Mark H Histed. Excitation creates a distributed pattern of cortical suppression due to varied recurrent input. Neuron, 111(24):4086–4101, 2023.

[39] Alfonso Renart, Jaime De La Rocha, Peter Bartho, Liad Hollender, Néstor Parga, Alex Reyes, and Kenneth D Harris. The asynchronous state in cortical circuits. science, 327(5965):587–590, 2010.

[40] Carl van Vreeswijk and Haim Sompolinsky. Chaotic balanced state in a model of cortical circuits. Neural computation, 10(6):1321–1371, 1998.

[41] Michael Okun, Nicholas A. Steinmetz, Lee Cossell, M. Florencia Iacaruso, Ho Ko, Péter Barthó, Tirin Moore, Sonja B. Hofer, Thomas D. Mrsic-Flogel, Matteo Carandini, and Kenneth D. Harris. Diverse coupling of neurons to populations in sensory cortex. Nature, 521(7553):511–515, May 2015. ISSN 1476-4687. doi: 10.1038/nature14273. URL https://doi.org/10.1038/nature14273.

[42] Douglas A Ruff, Joshua J Alberts, and Marlene R Cohen. Relating normalization to neuronal populations across cortical areas. Journal of Neurophysiology, 116(3):1375–1386, 2016.

[43] Kenneth D Harris and Thomas D Mrsic-Flogel. Cortical connectivity and sensory coding. Nature, 503(7474):51–58, 2013.

[44] Tania A Seabrook, Timothy J Burbridge, Michael C Crair, and Andrew D Huberman. Architecture, function, and assembly of the mouse visual system. Annual review of neuroscience, 40(1):499–538, 2017.

[45] Tobe CB Freeman, Séverine Durand, Daniel C Kiper, and Matteo Carandini. Suppression without inhibition in visual cortex. Neuron, 35(4):759–771, 2002.

[46] Steffen Katzner, Laura Busse, and Matteo Carandini. Gabaa inhibition controls response gain in visual cortex. Journal of Neuroscience, 31(16):5931–5941, 2011.

[47] Joonyeol Lee and John HR Maunsell. A normalization model of attentional modulation of single unit responses. PloS one, 4(2):e4651, 2009.

[48] Amy M Ni and John HR Maunsell. Neuronal effects of spatial and feature attention differ due to normalization. Journal of Neuroscience, 39(28):5493–5505, 2019.

[49] Ruben Coen-Cagli and Selina S Solomon. Relating divisive normalization to neuronal response variability. Journal of Neuroscience, 39(37):7344–7356, 2019.

[50] Oren Weiss, Hayley A Bounds, Hillel Adesnik, and Ruben Coen-Cagli. Modeling the diverse effects of divisive normalization on noise correlations. PLOS Computational Biology, 19(11):e1011667, 2023.

[51] Jonathan W Peirce. Psychopy—psychophysics software in python. Journal of neuroscience methods, 162(1-2): 8–13, 2007.

[52] Thomas A Pologruto, Bernardo L Sabatini, and Karel Svoboda. Scanimage: flexible software for operating laser scanning microscopes. Biomedical engineering online, 2:1–9, 2003.

[53] Marius Pachitariu, Carsen Stringer, Sylvia Schröder, Mario Dipoppa, L Federico Rossi, Matteo Carandini, and Kenneth D Harris. Suite2p: beyond 10,000 neurons with standard two-photon microscopy. BioRxiv, page 061507, 2016.

[54] Pauli Virtanen, Ralf Gommers, Travis E Oliphant, Matt Haberland, Tyler Reddy, David Cournapeau, Evgeni Burovski, Pearu Peterson, Warren Weckesser, Jonathan Bright, et al. Scipy 1.0: fundamental algorithms for scientific computing in python. Nature methods, 17(3):261–272, 2020.

[55] Charles R Harris, K Jarrod Millman, Stéfan J Van Der Walt, Ralf Gommers, Pauli Virtanen, David Cournapeau, Eric Wieser, Julian Taylor, Sebastian Berg, Nathaniel J Smith, et al. Array programming with numpy. nature, 585(7825):357–362, 2020.

[56] Kael Dai, Sergey L Gratiy, Yazan N Billeh, Richard Xu, Binghuang Cai, Nicholas Cain, Atle E Rimehaug, Alexander J Stasik, Gaute T Einevoll, Stefan Mihalas, et al. Brain modeling toolkit: An open source software suite for multiscale modeling of brain circuits. PLOS Computational Biology, 16(11):e1008386, 2020.

[57] Matthias Kaschube, Michael Schnabel, Siegrid Löwel, David M Coppola, Leonard E White, and Fred Wolf. Universality in the evolution of orientation columns in the visual cortex. science, 330(6007):1113–1116, 2010.

